# Livestock market data for modeling disease spread among US cattle

**DOI:** 10.1101/021980

**Authors:** Ian T. Carroll, Shweta Bansal

**Affiliations:** Department of Biology, Georgetown University, Washington, DC, USA; Fogarty International Center, National Institutes of Health, Bethesda, MD, USA

## Abstract

Transportation of livestock carries the risk of spreading foreign animal diseases, leading to costly public and private sector expenditures on disease containment and eradication. Livestock movement tracing systems in Europe, Australia and Japan have allowed epidemiologists to model the risks engendered by transportation of live animals and prepare responses designed to protect the livestock industry. Within the US, data on livestock movement is not sufficient for direct parameterization of models for disease spread, but network models that assimilate limited data provide a path forward in model development to inform preparedness for disease outbreaks in the US. Here, we develop a novel data stream, the information publicly reported by US livestock markets on the origin of cattle consigned at live auctions, and demonstrate the potential for estimating a national-scale network model of cattle movement. By aggregating auction reports generated weekly at markets in several states, including some archived reports spanning several years, we obtain a market-oriented sample of edges from the dynamic cattle transportation network in the US. We first propose a sampling framework that allows inference about shipments originating from operations not explicitly sampled and consigned at non-reporting livestock markets in the US, and we report key predictors that are influential in extrapolating beyond our opportunistic sample. As a demonstration of the utility gained from the data and fitted parameters, we model the critical role of market biosecurity procedures in the context of a spatially homogeneous but temporally dynamic representation of cattle movements following an introduction of a foreign animal disease. We conclude that auction market data fills critical gaps in our ability to model intrastate cattle movement for infectious disease dynamics, particularly with an ability to addresses the capacity of markets to amplify or control a livestock disease outbreak.

**Author Summary:** We have automated the collection of previously unavailable cattle movement data, allowing us to aggregate details on the origins of cattle sold at live-auction markets in the US. Using our novel dataset, we demonstrate potential to infer a complete dynamic transportation network that would drive disease transmission in models of potential US livestock epidemics.

## Introduction

Livestock operations within the United States (US) must be vigilant against trans-boundary animal diseases, including the critical threat to agriculture posed by a re-introduction of foot-and-mouth disease (FMD) [1]. Livestock movements were critical to the experience of the United Kingdom (UK) in 2001, where FMD was also non-endemic and where measures to control its outbreak included a temporary, nation-wide ban on the transportation of any livestock [2]. During the so-called “silent spread” period before the movement ban, nine out of the 12 geographic clusters of the epidemic had already been exposed to the disease through movement of infected animals between livestock operations [3]. Eventual costs to the agricultural sector reached £3 billion, and 5% of the UK’s 11 million cattle were culled to control the disease [2,4]. In the US, potential impacts of livestock movement on an FMD outbreak among the 90 million cattle maintained for beef and dairy production [5] are not easily estimated, but one study with disease spread constrained to California’s 5 million beef and dairy cattle predicts economic losses in the tens of billions of dollars with hundreds of thousands of culls [6].

The spread of disease between livestock operations through movement of infected animals is an inherent risk of modern systems of meat and dairy production in the US [7]. Many factors influence the progression of cattle from widely dispersed calving operations to relatively few packing facilities, including long term trends in productivity and consumer demand, a complex decadal-scale cycle in cattle inventories, and biologically driven seasonal cycles [8,9]. Variation in the volume of livestock in transit can be epidemiologically significant: for example, one factor amplifying the 2001 epidemic of FMD in the UK was the timing of the outbreak, which coincided with a seasonal peak in the number of sheep moving through livestock markets [3]. The number of animals moved between operations doubly affects the consequences of an outbreak, once through direct increases in opportunities for disease transmission and again for any additional time involved in tracing the movements of infected animals. Delays in the discovery of infected premises, on the order of hours rather than days for a disease as virulent as FMD, are certain to impact epidemic size and economic costs [6].

To understand the implications of livestock movement on the potential for disease spread in the US, it is necessary to consider the options available to livestock producers who buy and sell cattle. Auction markets (also referred to as stockyards and sale barns) are the predominate method of sale for producers selling cattle and calves in the US, although privately negotiated sales, packer-owned feedlots and other alternatives are common [10]. Auction market sales involve the movement of cattle to a stockyard, where they are grouped into lots, sold and transported to various buyers; alternative marketing methods avoid stockyards and vertically coordinate production chains [11], reducing the potential for contact between susceptible and infectious livestock. Producers selling through auction markets additionally influence the potential spread of a disease through choice in marketing areas, which tend to be large and overlapping [12, 13]. Despite extensive economic theory on when, where and how producers ought to market cattle, no data on real movements to or from auction markets have been collected to inform epidemiological analysis in the US.

FMD preparedness warrants close attention to auction markets, which played a central role in early, rapid expansion of the 2001 FMD epidemic in the UK [14] and possibly in several other countries with recent outbreaks of non-endemic FMD [15]. Epidemiological contact rates with auction markets in parts of Colorado and Kansas [16], California [17], and Texas [18] have been estimated through industry participant surveys. The most recent national survey indicates that over half of beef producers sold non-breeding stock at auction markets in 2006-07 (60.7% of producers for steers, 58.3% for cows [19]). Dairy contributes fewer US cattle shipments, but cows removed from dairy operations are also predominantly marketed at a stockyard [20]. While such surveys indicate the volume of cattle moved through particular marketing areas, they do not resolve movements between specific buyer or seller locations and individual stockyards, the kind of data increasingly in demand for emerging, epidemiological models for the spread of livestock disease [21].

Network models provide a layer of abstraction between raw data on individual contact rates, here cattle movements, and disease models [22]. Their use is primarily motivated by heterogeneity in the number of disease transmitting contacts attributed to infectious nodes (i.e. individuals or sub-populations), a pattern that emerged strongly during the initial spread of FMD in the UK’s 2001 epidemic [14]. Transportation networks for several European livestock industries [23–28] are known because animal tracking systems are mandated by the European Parliament [29]. Networks inferred from these data have lead to several results on surveillance and control strategies; for example, (1) identification of “sentinel” livestock premises projected to become infected early during an outbreak in Italy [30], (2) validation of risk reduction from the standstill rules implemented in the UK after 2001 [31], and (3) evaluation of targeted movement bans that selectively eliminate high risk contacts [28]. Network models for the UK cattle production system have additionally provided a foundation for livestock transportation strategies that promise efficient control of *endemic* diseases [32].

Livestock movement networks in the US have previously been inferred from interstate certificates of veterinary inspection (ICVIs), which provide the only regularly-collected record of cattle movement in the US [33–35]. However, ICVIs are only reported for interstate movement when only 20% of shipments onto US beef operations originate over 100 miles away [36]. Additionally, ICVIs are not required under federal rules for cattle moved interstate to approved livestock marketing facilities (including auction markets, which can play an outsized role in disease spread) when their next movement is intrastate [37].

Here, we introduce a novel stream of data on livestock movement through auction markets in the US, summarize the record of actual movements collected to date, and implement an approach to fitting a dynamic network model for livestock movement to this dataset. The data source is auction markets themselves, some of which publicly share the location of origin for representative lots of cattle sold on a particular day as advertisement for their business. Estimating a network model requires two levels of extrapolation from these data. First, we demonstrate that a sampling model allowing for strong overdispersion extrapolates locations of origin among representative lots to locations for all cattle received at auction. Second, we relate the patterns of movement to spatial covariates from the agricultural census, demonstrating potential to infer unobserved movements of cattle to auction markets that do not issue public market reports. We conclude with a model implementing the spread of disease on a livestock movement network; the model demonstrates how market biosecurity practices impact epidemic prevention in the aftermath of a foreign animal disease introduction. Animal tracing rules promulgated by the US Department of Agriculture (USDA) [37] forestall detailed movement tracking for predictive disease modeling, so we anticipate that our data mining approach will continue to be the only avenue for recording actual, intrastate and auction market movements of cattle in the US.

## Methods

As an integral part of the livestock system, auction markets are distributed through-out the US (Box 1A) [38]. USDA market reporters attend sales (i.e. the live auction of multiple lots of cattle occurring on a particular day) for many auction markets in order to collect volume and price information that is distributed through the USDA Agricultural Marketing Service’s Market News (MN). To provide additional information for advertising purposes, market reports may be published on websites maintained by individual auction markets. These independent market reports include one or more of: the total number of cattle sold which is referred to as “receipts”, price ranges for different types of cattle, and the winning bid for particular “representative lots” of cattle. Descriptions of representative lots include the source of cattle, the type and number of cattle and the price obtained. The total number of cattle within representative lots rarely approaches receipts in any market report, so both receipts and representative lots are relevant to estimation of cattle movement networks.

**Box 1.**
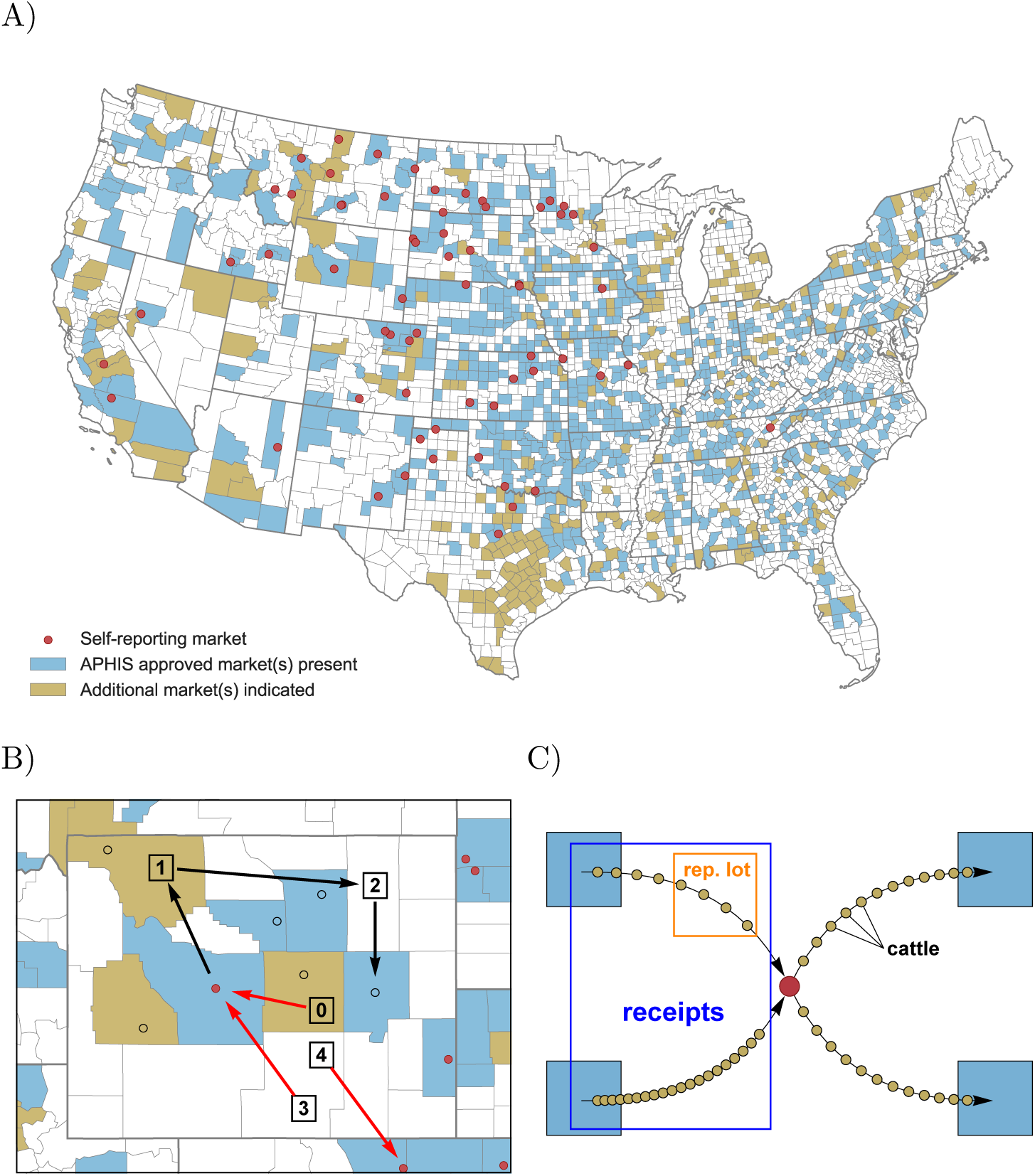
Cattle sales at auction markets are an integral part of the US livestock system. Marketing facilities, in-particular those voluntarily approved by APHIS and subject to ICVI exemptions, are distributed widely throughout the US. Representative lots of cattle sold at markets with online market reports provide a sample of the cattle transportation network. A) Presence of at least one auction market in each US county (blue or gold), or at east one APHIS-approved marketing facility (blue), estimated by compilation of public and private market directories [38]. Auction markets reporting representative lots collected for this study (red points) are concentrated in major beef producing regions. B) Two auction markets in Wyoming advertise representative lots online, revealing movement of cattle from unspecified livestock operations within a known county (numbered squares) to those markets (red arrows). Black arrows represent movement of cattle that is unobserved because the cattle are leaving a market, consigned at a non-reporting market (black circles), or moved directly between livestock producers. Individual animals may pass through multiple markets and farms; for example, a cow-calf operation (0) consigns a weaned steer calf at auction that is purchased by a stocker/back-grounder (1), who has a marketing agreement with a feedlot (2), that consigns the fed steer to an auction market buyer for a meat packing operation (not shown). Our data processing aggregates farms within counties, so farms 3 and 4 are represented in a movement network by a single county node connected to two market nodes. C) Representative lots are small groups of cattle consigned by one operation on the day of sale at a given auction, and receipts are the total number of cattle consigned at that sale. Purchased lots are typically combined with others, but could also be disaggregated, for shipment to the buyer’s operation(s).

We define the epidemiologically-relevant contact network as a bipartite network between county nodes (containing multiple farms) and market nodes, connected by edges representing movement of cattle between a county (or the farms in it) and a given auction market (Box 1B). Our data result from egocentric sampling on the market nodes. Receipts correspond to the rate of flow (i.e. node strength) through a given market node; and representative lots represent a sample of movement (i.e. edge weights) from origin to destination nodes (Box 1C).

In the first section below, we describe the aggregation of data compiled from market reports across market-owned websites. The following section explains methods for estimation and validation of a network model, built on the newly collected data and for the purpose of driving national-scale cattle disease simulations. These methods lay the groundwork for a network model serving to extrapolate from our data to a complete network of market-directed cattle movement in the US. Data on receipts and representative lots collected for this study are freely available online (Supplementary Data).

### Data collection

We automated the aggregation of receipts and representative lots across livestock auctions held at various auction markets in the US (Box 1A). Overlapping and incomplete directories of auction markets are maintained by multiple regulatory agencies and one business association. We compiled four such directories to identify auction markets [38] with potential to publish representative lots. The Livestock Marketing Association [39] uniquely provides websites of auction markets, so we manually searched the 322 websites included in the LMA directory for market reports. We wrote software in Python to download and parse information from websites that regularly (usually weekly) publish market reports including representative lots and which permit crawling by the Robots Exclusion Protocol. For each market report, the software cross-references the date against the market’s corresponding series of reports available through the MN search [40] to relate our records to a MN report on the same sale, when available.

For each representative lot in a market report, the software parses the consignor’s (i.e. seller’s) location, cattle type (e.g. steer or heifer), number (or “head”) of cattle, their average weight, and the purchase price (either per head or per hundred weight). It also extracts total receipts from any market report that includes it as well as receipts from the corresponding MN report if available. Where necessary, market reports are converted to plain text format from images with Tesseract OCR [41] or from PDFs with pdftotext [42]. A single animal per lot is assumed whenever the number of cattle cannot be parsed. We tuned the parser for each website until two researchers confirmed 100% accuracy of representative lots in at least two current market reports and a haphazard sample of archived reports, which varied widely in availability. Websites continue to be checked twice weekly for new reports, and parsers failing to obtain weekly data, returning data of the wrong type (i.e. string or numeric), or returning a head total from representative lots that exceeds receipts are manually checked and promptly corrected. The present analyses incorporate market reports obtained from June 2014 through Feb 28, 2016, including archived reports on sales dating from the first week of 2012.

The location of origin for a representative lot is typically given as a city or other populated place, with or without a state, and can be ambiguous. On the assumption that livestock operations are proximal to, but rarely within, the given city, our software georeferences each city to a US county, parish or equivalent (hereafter “county”). This loss of geographic precision is primarily a trade-off for accuracy, but the county scale also aligns with administrative boundaries relevant to disease management. The software matches locations, with common abbreviations expanded as needed, against names of populated places in North America using the GeoNames web-service [43] in order to identify the encompassing county. The county closest to the reporting market’s county, as determined by the great-circle distance (GCD) between county centroids [44], is recorded as the true location of origin. Our analyses include any market report with at least one representative lot whose origin could be georeferenced to a county by this procedure.

In order to extrapolate from collected market reports to unobserved movements of cattle, we incorporate cattle and calf inventory and sales from county tables in the USDA National Agricultural Statistics Service’s Census of Agriculture [5] as independent variables. After eliminating one of any pair of variables with strong correlation apparent on logged or untransformed axes, the following independent variables for each county remain: inventory of beef cows (*IB*), inventory of milk cows (*IM*), inventory of cattle on feed (*IF*), and sales by number of cattle over 500lbs (*SC*). Data for counties with few operations is often redacted, but we filled in most of the missing data by 1) summing over the product of the state average value per operation with the number of operations in the county for each operation size category, and then 2) subtracting non-redacted components from aggregated fields (e.g. inferring milk cow inventories as the difference between total cow and beef cow inventories). We created binary variables (e.g. *nzIB* for non-zero inventory of beef cows) to distinguish zero from non-zero values within each numeric variable and transformed the latter by taking z-scores of the logarithm of non-zero values (e.g. *lnIB*). A final group of variables for each county and market pair describes whether the county and market are in the same state (*S*), whether the market is in the county (*C*), the z-score of the non-zero distance between centroids of the county or origin and the market’s county (*D*), and the z-score of the logarithm of this distance (*lnD*).

### Inferring unobserved movements

We define our network model as a joint probability distribution over the edges between pairs of nodes, representing counties and auction markets. Each edge corresponds to the number of cattle from a given county (of *c* total counties) that are sold at a given auction market (of *m* total markets). A realization of the network model is the set of non-negative integers *ni*,*j*,*t* representing movement of cattle to market *i* from county *j* at each time period *t* (i.e. the total number of cattle on *one* of the left-hand edges in Box 1C). For each realization, the corresponding value for receipts consigned at market *i* at time *t* is 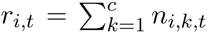. We employ a joint probability distribution over network edges of the form

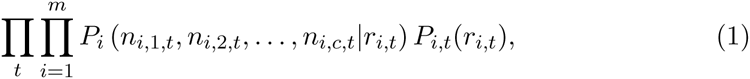

which embeds several simplifying assumptions. The foremost assumption is inde-pendence, both between markets and over time, for all edges representing cattle movements. Another implied assumption is that time-dependence only factors into the distribution on receipts, *P*_*i*,*t*_(·), or the total number of cattle sold at market *i* during period *t*. In other words, the number of cattle from each county has a constant probability distribution conditioned on receipts in this model. Choosing a model of this form facilitates estimation of two distinct factors, *P_i_*(· |·) and *P*_*i*,*t*_(·). We employ the first as a likelihood over representative lots and the second as a likelihood over receipts. We fit parameters of the likelihood models, described next, by sampling posterior parameter vectors with Hamiltonian Monte Carlo implemented in Stan [45] with weakly-informative Gaussian or half-Cauchy priors (Supplementary Methods).

The multivariate distribution on the number of cattle from each county, *P_i_*(· |·), has the sample space of a multinomial distribution, but the sampling method dictates a different likelihood. The variable represents a sum over the number of cattle in one or more representative lots from each county, and not, as in a multinomial distribution, a sum over independent trials with equal probability. The Pólya (a.k.a Dirichlet-multinomial) distribution, parameterized by non-negative *α*_*i*, ·_, flexibly admits the overdispersion resulting from this “bursty” sampling method [46]. We initially explored the suitability of using the Pólya family in our likelihood function by directly estimating *α*_*i*, ·_ for each auction market independently, i.e. including only the *i*^*th*^ market’s representative lots in the likelihood. Subsequently, we estimate posterior distributions over parameter vector *β_x_* affecting all markets and counties simultaneously through linear combination with independent variables *x*_*i, j*_. In this case, the first factor in the network model (1) has definition

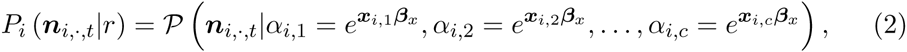

where *P* is the Pólya distribution’s probability mass function. Note that for model fitting, we take *n*_*i*,*j*,*t*_ to be a number of cattle observed in representative lots, for which the sum across counties is just the total head in representative lots and not the market’s receipts for that auction. Only when using (1) predictively do we take *n*_*i*,*j*,*t*_ to equal all receipts from the *j*^*th*^ county. The independent variables in *x*_*i*,*j*_ have fixed effects *D*_*i*,*j*_, *lnD*_*i*,*j*_, *S*_*i*,*j*_, *C*_*i*,*j*_, *nzIB_j_*, *nzIM_j_*, *nzIF_j_*, *nzSC_j_*, *InlB_j_*, *InlM_j_*, *InlF_j_*, *InSC_j_*, and a random effect of market, *M_i_* ~ *N*(0, *σ_x_*). Posterior predictive modeling, using MCMC samples of (*γ_x_*, is possible for any market for which *x*_*i*,*j*_ are available, which admits non-reporting markets in any censused county.

The network model (1) gains temporal dynamics by allowing the distribution on receipts, *P*_*i*,*t*_(·), to change over time. The volume of cattle traded at specific auction markets is driven by social and economic variables at several spatial and temporal scales, so we do not presently attempt to model receipts at auctions for which we did not obtain a market report. Instead, we model receipts for auction markets in relation to the number of cattle from all representative lots in a given market report, on the assumption that this total follows the trends in receipts. For receipts *r*_*i*,*t*_ at market *i* having representative lots totaling *s*_*i*,*t*_ head during time period *t*, we assume a negative-binomial distribution on the unobserved head, *r*_*i*,*t*_ – *s*_*i*,*t*_. Parameterizing its probability mass function by the mean *μ* and a parameter *ϕ* such that the variance is *μ* + *μ*^2^/*ϕ*, we set the second factor in the network model (1) to

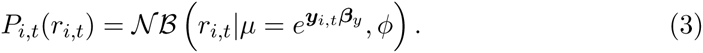

Posterior samples for (*β_y_* and *ϕ* are obtained using the subset of market reports for which we obtained receipts. The independent variables in *y*_*i*,*t*_ represent fixed effects for year (*Y_t_*) and week of year (*W_t_*) along with *s*_*i*,*t*_, *lnIB_i_*, *nzIM_i_*, *nzIF_i_*, *InlM_i_*, *InlF_i_*, *InSC_i_*, and a random effect of market with variance 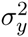.

### Validation

Markets contributing to the data stream are self-selecting, prompting the need for validation against randomly sampled data where possible. In addition, the number of cattle transported to a given auction from each county is a high-dimensional response variable with potential to violate the assumptions of our multivariate regression model. We validate the likelihood function assumed in our Bayesian inference by checking self-consistency: we test the goodness-of-fit between posterior predictive distributions and observed data. Specifically, we estimate posterior predictive distributions for two summaries of the data, the number of counties represented at a given market averaged across market reports and the average distance cattle are moved to a given market, and compare to empirical distributions using the one-sample KS test. To seek evidence against biases in the underlying data itself, we compare our movement data to previous estimates of livestock transportation. Freely available data on livestock movements in the US is only available for year 2001 health-certificate derived data on interstate flows [7], but these data have recently been validated against ICVIs from year 2009 [33]. We aggregated all representative lots from a full year of market reports by the markets’ state, and correlated the number of cattle by state of origin with year 2001 interstate flows.

## Results

We report on sales from Jan 2, 2012 to Feb 28, 2016, with a sample size of 13 markets in 2012 increasing to 63 markets by 2016 (Fig. 1A). Cattle from representative lots in 5171 reports from 63 markets located in 60 counties in 17 states are represented in this analysis (Box 1A). The total number of cattle included among 347 thousand representative lots is 3.1 million. In the three sections which follow, we summarize overall trends observed in the data aggregated across market reports, show the predictability of the county of origin for cattle sold at different markets, and relate the number of unobserved cattle sold at a given auction to both static and time-varying covariates.

**Figure 1.**
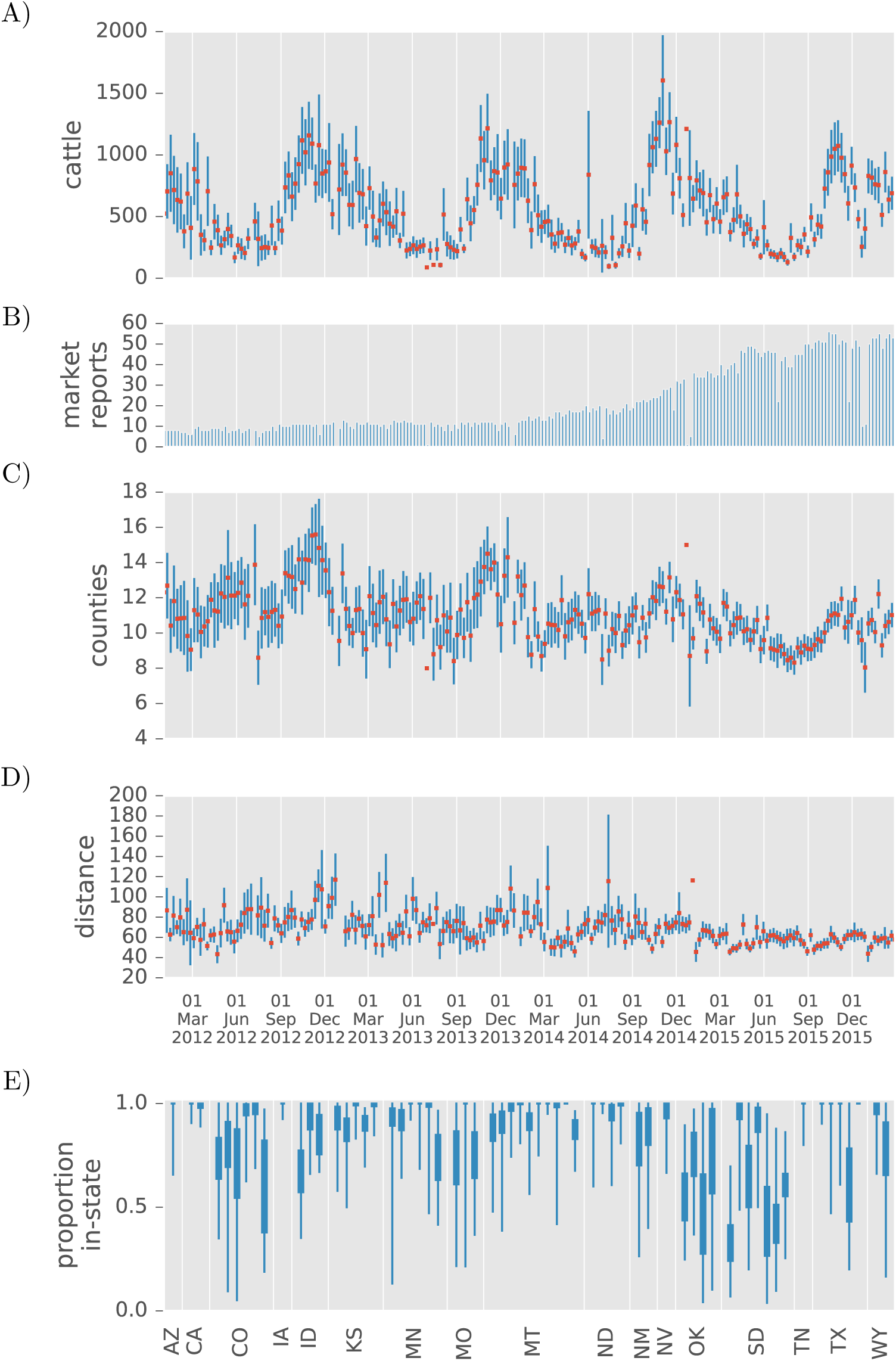
Cattle movements captured in representative lots for a 3.5 year time span from auction markets in the US. A) The total number of reports collected for cattle sales occurring within a given week. B) The average (+/- SEM) head of cattle reported with locations of origin each week by a single livestock market. The outlying observation in the first week of June, 2014 includes a massive auction conducted by the World Livestock Auctioneer Champion in Ft. Pierre, SD. C) The average (+/- SEM) number of counties from which cattle arrived for sale at a market on a given week. D) The average (+/-SEM) distance between county of origin and auction market per head of cattle sold on a given week. E) The proportion of cattle (interquartile and entire ranges) in representative lots transported from a county within the same state to each reporting market.

### Static and seasonal patterns

The average number of cattle movements reported in representative sales for a live auction increases to a peak of 1000-1500 head in late fall and decreases to a few hundred during summer months (Fig. 1B). This seasonal trend persists for the duration of our study period. For a given week, livestock markets report receiving cattle from 11 (SD 1.5) counties on average, and this lower bound on network “in-degree” also shows seasonal variation with increases of a lesser magnitude in late autumn (Fig. 1C). Seasonal peaks for in-degree roughly follow the variation in cattle volumes, but grow less pronounced as the number of markets included in the dataset grows. Likewise, the number of counties represented during low seasons fails to track the volume of cattle before the most recent year, but the larger samples achieved since late 2015 show in-degree variation that is well correlated with variation in cattle volumes. The average distance between county of origin and auction market for cattle in representative lots is well under 100 miles in the vast majority of weeks (Fig. 1D), which s consistent with producer surveys [36].

Interstate movements are scarce among representative lots, but those observed are well correlated with pre-existing state in-shipment data. Aggregating across all markets, the proportion of sales that originate within-state shows no temporal trend in deviations from an average of 0.84 (SD 0.067). Oklahoma and South Dakota auction markets typically sold the most out-of-state cattle, and were the only states with a market that on-average advertised more out-of-state cattle than in-state cattle (Fig. 1E). Nonetheless, among the 16 states (Tennessee excluded) with year 2015 representative lots that could be compared to year 2001 in-shipments [7], the Pearson correlation coefficients for cattle by origin state are greater than 0.48 in all but two reliable estimates (Table 1). Representative lots marketed in Iowa, Arizona, and California were insufficient for meaningful comparison, and the lowest remaining coefficient (0.25 for Idaho) is driven by one strong connection to Nye County in Nevada. South Dakota has the largest number of interstate representative lots that we observed and has a strong correlation of 0.79. Montana and Colorado, while having similarly large sample sizes, exhibit opposite extremes with the first highly correlated at 0.90 and the other relatively weakly at 0.48.

**Table 1.** Correlations between interstate representative lots and inshipment data from Shields & Mathews [7]. For states with at least one reporting market, the table shows the Pearson correlation (*ρ*) for the number of cattle from *n* origin states, along with the number of cattle in the representative lots that moved between states and its proportion (*p*) of the total.

| Dest. | $p$ | Head | $n$ | $\rho$ |
| --- | --- | --- | --- | --- |
| IA | 0.99 | 20 | 34 | -0.04 |
| AZ | 0.99 | 25 | 15 | -0.10 |
| ID | 0.66 | 7174 | 43 | 0.25 |
| KS* | 0.95 | 5600 | 38 | 0.37 |
| CO* | 0.85 | 10915 | 23 | 0.48 |
| MO | 0.75 | 11544 | 17 | 0.49 |
| MN* | 0.96 | 4082 | 37 | 0.54 |
| CA | 1.00 | 19 | 43 | 0.66 |
| WY* | 0.86 | 10072 | 28 | 0.72 |
| OK | 0.68 | 6381 | 18 | 0.73 |
| NV | 0.97 | 166 | 28 | 0.77 |
| ND | 0.98 | 4300 | 3 | 0.78 |
| SD | 0.74 | 110644 | 39 | 0.78 |
| TX* | 0.98 | 287 | 33 | 0.79 |
| MT* | 0.92 | 16628 | 16 | 0.90 |
| NM | 0.83 | 4403 | 17 | 0.99 |

\* State import rules, as presented on each state’s agricultural department website, except certain livestock markets from ICVI requirements.

### Fitting to representative lots

A first test of the network model is whether any parameterization of the Pólya family can fit the high-dimensional data on the number of cattle by county for each market, leaving aside the regression against independent variables. The posterior mean of *α*_*i*,·_ (the Pólya parameters for market *i*) indicates the fitted strength of association between each market and different counties (Fig. 2A), but doesn’t directly correspond to the observed average number of cattle for that edge because of variation in the total number of cattle between market reports. A better visualization of the goodness-of-fit is the posterior predictive distribution on the number of counties represented as a function of this total (Fig. 2B), which falls below what would be expected under a multinomial sampling model whenever cattle from the same county are clustered in lots. These fitted curves capture the trend and variability in the county-distribution of representative lots for all but a handful of markets with smaller sample sizes (Supplementary Figures).

**Figure 2.**
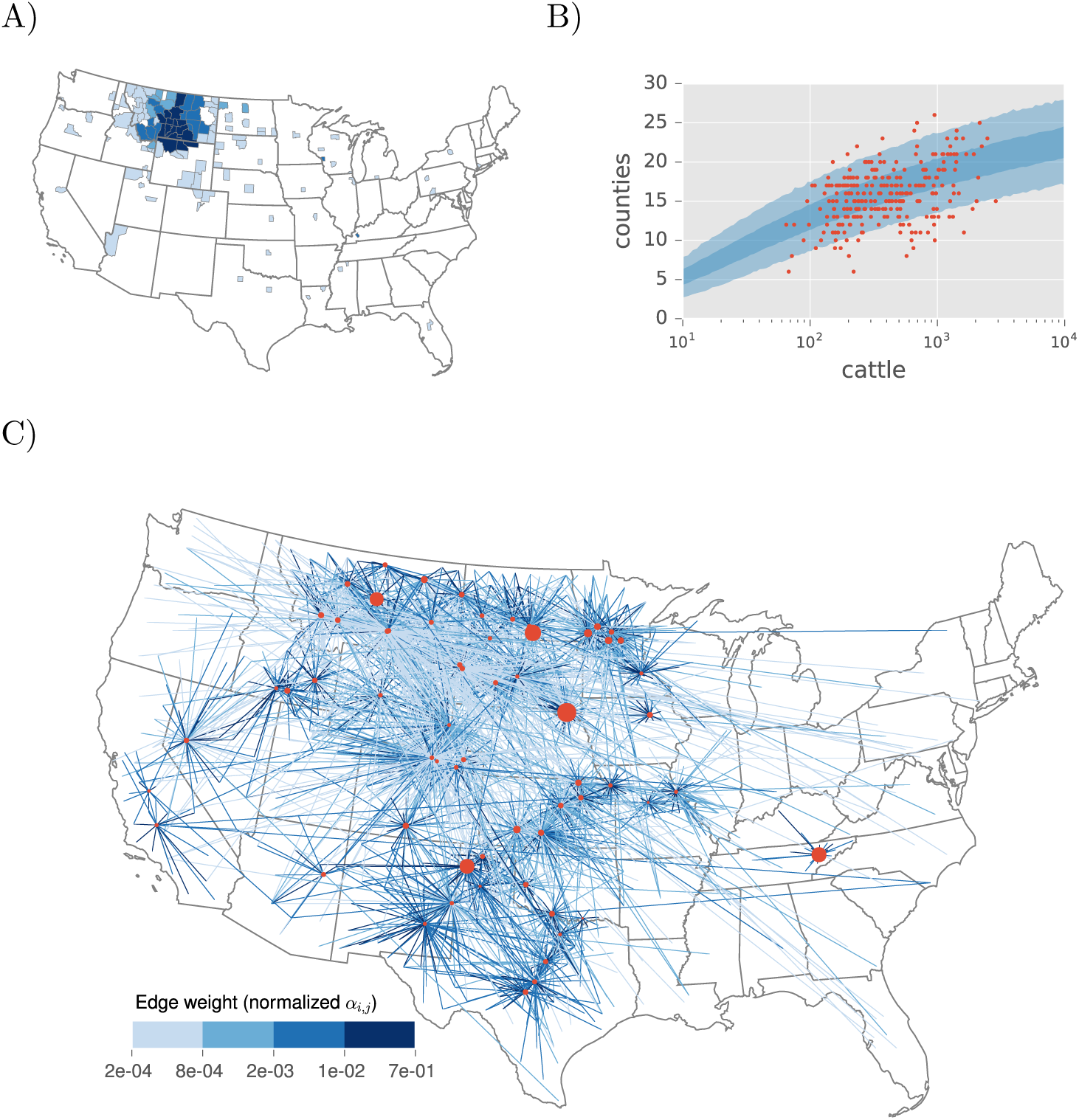
Values of the network model parameters, *α*_*i*,*j*_, that affect the distribution of counties from which cattle are sold when these are fitted directly and independently for each market. Panels A and B exemplify the fit obtained for nearly all markets (Supplementary Figures). A) The posterior mean *α*_*i*,*j*_ for a market *i* in Montana and counties *j* from this market’s representative lots. B) The predictive distribution (50th and 95th percentiles) on the same market’s in-degree for increasing receipts in comparison to individual market reports (points) showing the number of counties and total head within representative lots. C) The posterior mean *α*_*i*,*j*_ for all markets, rescaled as an edge weight represented by color, along with the log of the total *α*_*i*,*j*_ over all counties *j* represented as market vertex size.

Markets naturally exhibit variation in the weight of edges connecting to each county, quantified by the posterior mean of the normalized *α*_*i*,·_, as well as variation in the degree to which representative lots approach independent samples across counties, measured by the sum of *α*_*i*,*j*_ over *j* (Fig. 2C). Mapping the edges visualizes the sharp decline in weight with greater distance, anticipating a strong effect of distance when the *α*_*i*,*j*_ are related to independent variables. Variation among markets in the total of *α*_*i*,·_ indicates differences in overdispersion relative to a multinomial distribution, with a smaller total indicating more “bursty” arrivals of cattle from each county. Only a handful of markets have relatively large totals, indicating that most counties with farms consigning cattle to these few markets are represented at every sale. In the majority of markets, counties are more irregularly represented at a given sale, but show up with larger numbers of cattle when present. Another visually apparent distinction between markets is on the distribution of weights among edges pointing to a given market: the market in Nevada, for example, has a long tail of infrequently represented counties when compared to the market in Arizona.

In the second round of fitting to representative lots, using likelihood (2) to incorporate all representative lots conditioned on the total per market report, we predict edge weight and node strength with independent variables related to geography and agricultural production. Samples from the posterior distribution on *β_x_* are negative for both linear and logarithmic functions of distance (*D* 95% CI-1.90 to -1.86, *lnD* 95% CI -1.313 to -1.306) as anticipated. The county in which a market resides is typically well represented at auctions there (95% CI 10.6 to 10.7), and administrative boundaries also amplify expected movements to in-state markets (95% CI 0.76 to 0.78). Agricultural census data included in the model have different effects based on the class of cattle. The presence and abundance of cows in a county increased its edge weights (*nzIM* 95% CI 0.11 to 0.13, *IM* 95% CI 0.066 to 0.071, *nzIB* 95% CI 3.1 to 3.3, *IB* 95% CI 1.00 to 1.02), with beef cows having the larger effect. In contrast, the presence and abundance of cattle that are further along the chain of production (e.g. feeder cattle) reduced edge weights (*nzIF* 95% CI -0.20 to -0.18, *IF* 95% CI -0.084 to -0.075, *nzSC* 95% CI -4.01 to -3.88, *SC* 95% CI -0.35 to -0.34). We capture unexplained, between-market variation in the total of *α*_*i*,·_. with a random intercept (*σ_x_* 95% CI 1.07 to 1.15); however, the posterior mean coefficients indicate some pressure to deviate from the assumption of a normal distribution (Fig. 3A).

**Figure 3.**
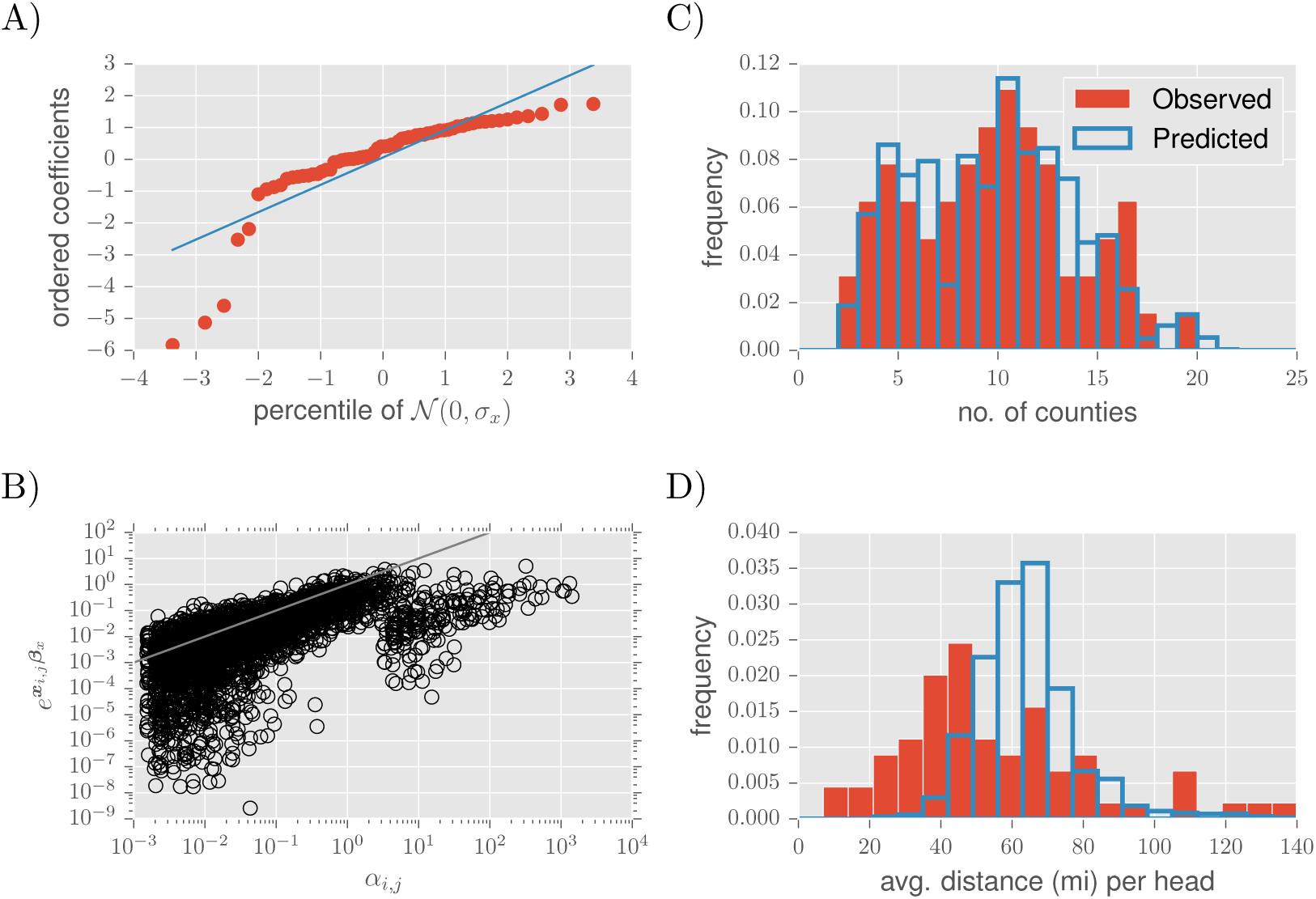
Network model parameters *β_x_* affect the distribution of counties sold at each market through linear combination with independent variables *x*_*i*,*j*_. A) Probability plot comparing the posterior mean market intercepts with a normal distribution having SD equal to the posterior mean of *σ_x_*. B) Comparison between the regression model and the saturated model, in which each *α*_*i*,*j*_ is a free parameter. C) The distribution on market average in-degree, or the average number of counties of origin among representative lots for a given auction market. D) The distribution on market average movement distance per head, the distance between centroids of the origin county and the market’s county averaged across animals in each market report and subsequently across reports for each market.

A comparison between observed data and posterior predictive distributions indicates both strengths and weaknesses in the regression model. The in-degree distribution across markets (averaged over market reports) does not deviate from the distribution generated by the assumptions of the fitted likelihood model (KS statistic 0.064, *P* > 0.1). In other words, given a total number of cattle in representative lots consigned at a given auction, the model groups them into the correct number of counties on average (Fig. 3C). However, the distribution across markets on the average distance cattle move to market deviates from the prediction (KS statistic 0.41, *P* < 0.001). There is greater variability among markets in the average distance that cattle are transported to market (Fig. 3D) than the regression model achieves, indicating the need for an interaction between distance-related covariates and individual markets.

### Fitting to receipts

The likelihood on receipts captures the dynamic nature of the movement network by positively responding to variation in the total number of cattle in representative lots and including non-zero coefficients for different weeks and years (e.g. Fig. 4A). Two distinct processes represented by these coefficients drive temporal variation in receipts within a market. On average, auction markets report representative lots in proportion to the receipts for a given sale, posting more representative lots when more cattle are sold (95% CI 0.47 to 0.49). This proportion, however, is not constant over time as additional increases in un-observed receipts are associated with certain weeks (e.g. 95% CI 0.36 to 0.52 in late October) and decreases occur in others (e.g. 95% CI -0.59 to -0.38 for late July). The time-varying fixed effects combine with the negative binomial overdispersion parameter (*ϕ* 95% CI 4.3 to 4.4) to produce realistic levels of within market variation in the proportion of receipts appearing in representative lots (Fig. 4B); the remaining independent variables, which also happen to be static, account for between market variation. The occasional underestimate of receipts during peak cattle trading season could likely be resolved by relating *ϕ* to some of these same independent variables.

**Figure 4.**
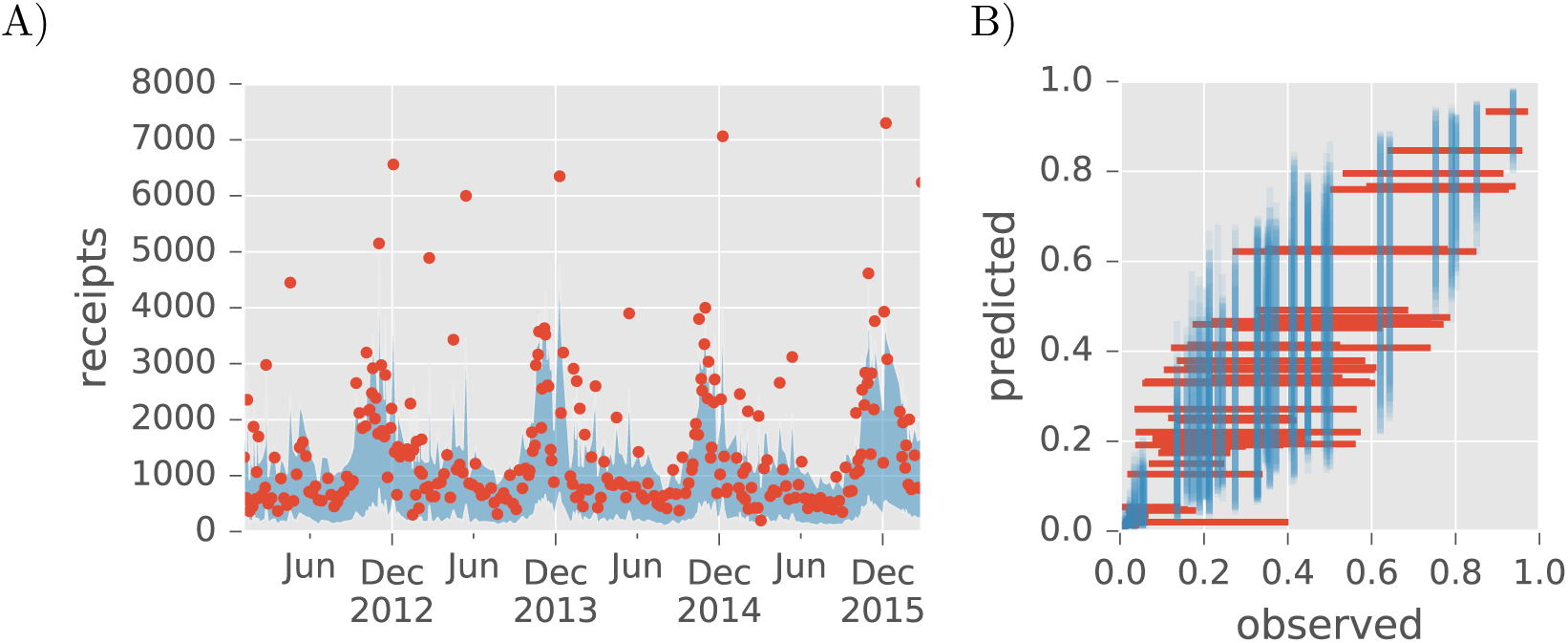
Network model parameters (*δ_y_* affect the number of receipts at each market through linear combination with independent variables *y*_*i*,*j*_. A) The posterior predictive range (95th percentile) on receipts for the Montana market from Fig. 2A with observed receipts (points). B) Within and between market variation in the proportion of receipts included among representative sales. For each market, lines show the inter-95%-tile range of observed (horizontal) and predicted (vertical) proportions intersecting at their respective means.

Agricultural census data for the county in which an auction market is located statistically explains part of the variation between markets, and the remainder is attributed to a random effect of market. Non-zero inventories of milk cows (95% CI -3.5 to -2.7) and cattle on feed (95% CI -2.5 to -1.4) both reduce the expected number of un-observed receipts, and the magnitude of cattle inventories or sales have further negative effects (*lnIM* 95% CI -0.45 to -0.35, *lnIB* 95% CI -0.79 to -0.55, *lnIF* 95% CI -0.83 to -0.65, *lnSC* 95% CI -0.40 to -0.086). Each market has a random intercept with standard deviation *σ_y_* (95% CI 0.68 to 0.73) that is large compared to the standard deviation among markets for the combined effect of the census variables (95% CI 0.010 to 0.12), implying that much of the variation in the unobserved receipts (Fig. 4B) is “explained” by unknown sources of variation between markets. In addition, for a handful of markets, the static census data for that county drives the predictive distribution for un-observed receipts well below the true values, suggesting that some critical process is not represented in the covariates (Supplementary Figures).

## Application: Modeling Disease Spread

To illustrate the utility of our network model, we study infection propagation from county to county, thwarted by market biosecurity measures. For this, we develop a simple infection spread model driven by variability in the number of cattle arriving at market from a county. In conjunction with movement bans for designated “control areas” [35] that serve as the primary strategy for containment of an FMD outbreak in the US [1], biosecurity procedures at auction markets are a critical mechanism for preventing movement of potentially infectious animals between livestock operations. The premise is that biosecurity practices at auction markets can provide a firewall, preventing infection from reaching new operations by intercepting cattle that may be carrying disease from an operation with infected animals. Our model tracks the number of counties (we assume that movement bans are implemented one whole county at a time) that must be designated as control areas during eradication of an FMD-like outbreak. All exports from a control area are restricted, but movements that occur before the control is established are responsible for growth in the number of counties affected. The way an area becomes eligible for control is through a “dangerous contact”, which we define as receiving cattle from any farm in an area that previously had a dangerous contact or was infected. We assume here that all movements occur through a livestock market, and that contact between animals at a livestock market does not engender additional dangerous contacts between operations. In summary, a county becomes a control area if it receives a shipment of cattle from any county that is also destined to become a control area, but has not become one yet.

We model disease spread through a branching process [47 section 15.2] that extends *n* generations if spread is halted by the time any one county is at most *n* dangerous contacts removed from the county at the source of the outbreak. The model is completely specified by choosing a distribution on the number of distinct counties that could have a dangerous contact with a single county. We denote this random variable by *Y* and estimate its distribution using the above likelihood models, with parameters sampled from estimated posterior distributions given the market data, and additional assumptions (Supplementary Methods). Biosecurity practices at auction markets are incorporated into the distribution on *Y* as a zero-inflation parameter. With probability *τ*, a market quarantines cattle arriving from a farm within a county that will become a control area, resulting in no additional control areas becoming established due to consignments at that sale. The remaining parameters in our model may be explained through their effect on the expectation 〈*Y*〉 = *λ*(1–*τ*)*ν*〈*N*〉: *λ* is an average number of marketings, *ν* is a splitting parameter for the allocation of cattle from the origin county among destination counties, and 〈*N*〉 is the expected number of cattle per origin county per sale. Then, the expected value of the total number of counties, *X_n_*, that must become control areas in a branching process stopped after *n* generations, is

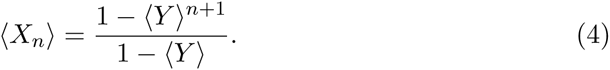

Additional details in the Supplementary Methods justify this result under the strong assumption that the distribution on *Y* is the same for any county which becomes a control area, which is only reasonable for small *n*.

Variation in the factors contributing to 〈*Y*〉 drive the effort needed to eradicate the disease; in particular, they determine where the critical threshold lies that separates a finite branching process from a potentially infinite one. The key contribution from a network model fitted to market report data is estimates of the number of cattle consigned at a given sale from farms within a single county. The estimate must be extrapolated from a combination of representative lots and receipts, which the fitted network model (1) rightfully predicts to be greater than direct observations (Fig. 5A). Temporal variation in the number of cattle marketed per county, which is slight but still apparent in both the observed and predicted distributions, is primarily due to variation in total cattle volumes (Fig. 1B). These volumes fluctuate more than the number of counties represented at auctions (Fig. 1C), which would otherwise have a counter-effect on the number marketed per county at different times of year. For fixed values of *λ* and *ν*, weekly variation over the course of the year in cattle sales strongly affects the threshold value of τ separating the two qualitatively distinct outcomes of the branching process (Fig. 5B). During more active cattle trading seasons, biosecurity must be considerably stronger to cause convergence of *X_n_* as *n* → ∞ or, equivalently, to bring the spreading process to a stop without further intervention. Averaging over the seasonal variation in cattle trade, for the purpose of considering introduction of an FMD-like disease at a randomly selected time of year, the effect of market biosecurity strongly influences the magnitude of any resulting eradication effort. The value of 〈*X_n_*〉 is the minimum number of control areas to establish if back-tracing to the origin from the county of detection involves a sequence of *n* dangerous contacts. Thus, interpreting *n* as a measure of the delay between disease introduction and a regional movement restriction that halts further dangerous contacts, we see that around the critical value of *τ*, doubling the delay nearly doubles the number of control areas needed (Fig. 5C).

**Figure 5.**
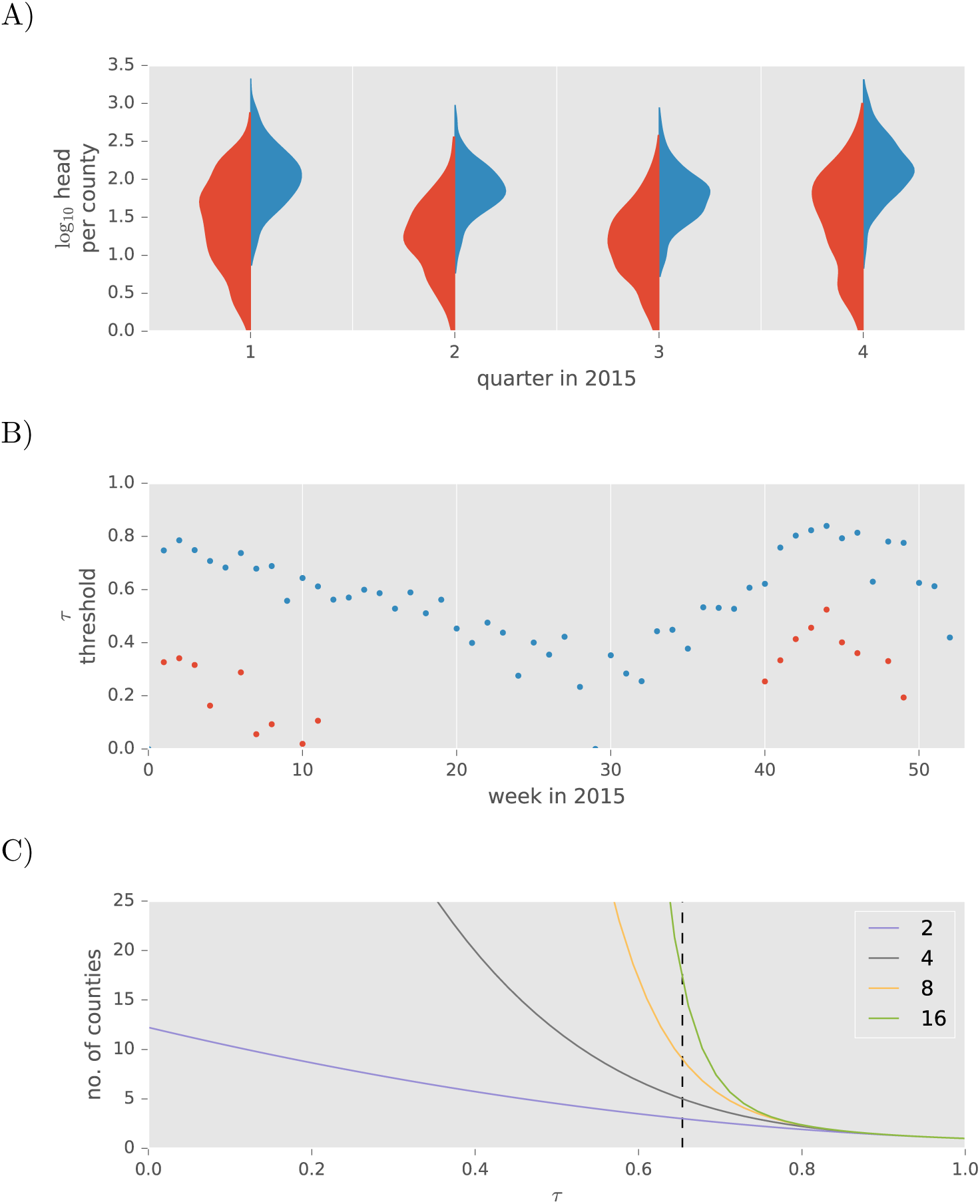
Influence of model parameters on the number of counties designated as control areas following a single introduction of a foreign disease of cattle. A) Observed (red) and modeled (blue) distributions on the the number of cattle shipped to any one market from any one county aggregated to quarters. B) The time-varying threshold for market biosecurity *τ*, that leads to a full-blown epidemic (blue). For comparison, we show results where the numbers of cattle involved in calculating the distribution on *Y* are estimated from raw averages across representative lots (red). C) The number of counties that become control areas across the range of *τ* for an outbreak contained after longer delays, interpreted as the maximum number of dangerous contacts a county must back-trace to reach the source of a disease outbreak. Parameter *λ* = 1 and *ν* = 0.02.

## Discussion

The epidemiological contact network is a fundamental component of models for the spread of a disease, and market reports publicized by livestock auction markets can contribute urgently needed data to support inference of such networks within the US livestock system. In this report, we describe an ongoing process to opportunistically sample cattle movements originating from counties with beef or dairy operations and consigned at auction markets distributed across the US. Our study complements previous efforts to summarize transportation of cattle within the US using data derived from interstate certificates of veterinary inspection [7,34], but is novel because it includes within-state movements and targets movements involving auction markets, which can be crucial to amplification or control of a foreign animal disease. We develop a statistical model for inference of a bipartite contact network between nodes representing either a US county (containing cattle farms) or a US cattle auction market, and demonstrate its link to epidemiological models of economically disruptive livestock diseases.

Data on cattle movements that are voluntarily reported by auction markets have not previously been aggregated and collectively analyzed, for either economic or epidemiological purposes. In contrast, data on volumes and prices from auction markets is central to the economic analysis of livestock systems, is naturally of keen interest to industry participants, and its reporting is mandated by law [48]. A few studies have examined data on movement of cattle sold through auctions [12,49], but those studies focus on internet auctions which do not imply movement of cattle through a physical stockyard. It is critically important to archive data released by auction markets while it is still available, and in particular before components of this novel data stream dry up. Recent declines in the reliance on auction markets for slaughter cattle have become severe enough to raise new concerns about price discovery, the backbone of an efficient beef market [48]. While such declines affect the economically important class of slaughter cattle, reliance on auction mar-kets for cattle with epidemiological significance, those moving between farms (e.g. replacements) or to feeding operations, shows no sign of decline (Supplementary Figures).

Cattle tracing systems in the UK and Europe create opportunities for research on a fully censused network, but the situation in the US is such that relevant network data must be sparsely sampled and produces a challenging population inference problem. In network sampling theory, randomly selecting nodes without tracing their edges to sample additional nodes, is an ego-centric sampling method that allows straightforward estimation of population level node attributes, but not of edge attributes [50]. In contrast, were edges randomly sampled as in the ICVI records, the reverse problem applies: the sample of nodes involved is automatically biased towards those of higher degree. Sampling theory pertaining to higher-order network properties, such as clustering and assortativity, becomes exceedingly complicated. Lacking a universally applicable sampling method for network data, what’s needed are computationally feasible methods for estimating the complete network in a manner that respects the likelihood of observing each network attribute with a given sampling strategy [51]. Our study lays the groundwork for extrapolating a complete network of cattle transported to auction markets from an ego-centric sample of representative lots and receipts at individual market nodes. While we might reasonably estimate average market degree (i.e. the number of connected counties) with a simple sample mean, we would not, for example, estimate the expected number of cattle moved on any edge in the complete network with a simple mean across representative lots. Instead, we propose a regression model with a flexible likelihood function to scale up from the known constraints of our sampling method. Our success at obtaining posterior probability distributions for the parameters relating cattle movement to independent variables is due to an efficient HMC sampler [45] in combination with an efficient algorithm for calculating a Pólya likelihood for multiple samples (Supplementary Methods). Incorporation of variables beyond the agricultural census data is computationally feasible in this framework, which already promises reliable estimates.

Prior to model fitting, we confirm that our data is generally consistent with known patterns and trends in the livestock industry. For example, seasonal variation in the volume of representative lots is consistent with beef cattle production systems, where calves are produced in spring and weaned cattle or yearlings sold to pasturing or feedlot operations in the fall and subsequent spring [52]. In addition, the average distance most cattle in representative lots move to a market is under 100 miles, while considerably longer distance movements do infrequently arise, as in survey results [36]. Likewise, the majority of observed movements are destined to a within-state market, which is consistent with the dominance of intrastate movements extrapolated by Lindström et al [33] from entirely interstate data. As these authors point out, the intrastate estimates could not be validated at the time; now, however, visual comparison between their state specific estimates [Fig. S4 in 33] and our data (Fig. 1E) indicates that market reports show even higher rates of intrastate movement. The un-matched data on cattle movements from market reports shows that roughly 80% of market directed shipments on any week originate within state. Among the remaining interstate movements, consignments from out-of-state counties correlate fair to strongly with state in-shipments summarized by Shields and Mathews [7]. Sale at livestock markets is not the only impetus for cattle transportation, but the correlation between representative lot origins and state in-shipments demonstrates its importance. Indeed, if the certificate-derived data do sample all movements (excepting slaughter animals) without bias, then the stronger correlations in Table 1 would indicate states for which cattle in-shipments are in fact movements to a livestock market. Alternatively, differences in ICVI requirements for cattle bound for livestock markets could affect the degree of correlation, but no such connection is apparent among the states with reporting markets.

Our confidence in the data quality extends also to the statistical model, which provides a strong starting point for predicting unobserved movements to non-reporting livestock movements. Opportunistic sampling unavoidably raises the specter of biased estimation on the population as a whole; any unknown associations between self-reporting market nodes and the volume of cattle they auction or the weight of edges from particular county nodes would bias network estimation. Regression of independent variables against receipts and representative lots incorporates some of these associations, eliminating a few potential sources of bias. More importantly, however, agreement between the sign of fitted coefficients and our intuition about the livestock system builds confidence in the model’s predictive ability. The negative effect of distance, for example, is one clear example of the model following intuition. The difference between the positive effect of cows and negative effects of feeder cattle on the weight of a county’s connections to markets was not anticipated, but is consistent with auction markets’ role in moving livestock from cow-calf operations towards feeding operations. The regression model for receipts naturally associates larger receipts with auctions for which a greater number of cattle are reported in representative lots, but it also indicates a downward adjustment in the predicted difference between receipts and total cattle in representative lots for markets in counties with larger inventories of cattle. The latter effect implies that such markets include a greater proportion of their receipts among the representative lots–a potentially rewarding strategy for markets advertising in active livestock producing regions. Our current regression models do not address all potential sources of bias in network estimation; for example, the markets sampled are predominately located in central states and any distinction between beef and dairy cows in the data has not yet been explored. Ideally, as for validation of any statistical model fitted to mined data, or otherwise opportunistic samples, we would compare model predictions to newly collected and truly random samples of the transportation network. In the context of cattle movements, however, the privacy interests of the livestock industry are a significant obstacle to collecting such validation data.

The central weaknesses of the combination of data and network estimation reported here is whats left out, movements of cattle leaving livestock markets and extension of the model for receipts to non-reporting markets. The model for receipts introduced here relies in part on information (the total head in representative lots) only available for auctions accompanied by market reports. Unlike the conditional distribution on edges (2) then, the predictive distribution on receipts cannot immediately extend to auctions from non-reporting markets. Estimating re-ceipts independently of partial data on a given auction requires a mechanistic model that takes into account seasonal and longer term dynamics driven by economic (e.g. prices of beef and feed) processes as well as reproductive cycles. Fortunately, there is a wealth of data on auction market receipts in the USDA Agricultural Marketing Service’s Market News archives to aid in model development. A greater challenge lies in estimating the markets’ out-going edges in the livestock transportation market. Information on the location or identity of cattle buyers is unlikely to be shared by any auction market, in order to protect their business. A more fruitful avenue for future research likely involves combining livestock market information with ICVI records, joining the data on both the location of origin (i.e. the market location) and the date of sale.

Auction markets can play a significant role in amplification of livestock disease, just as airports play a role for human diseases. The market report data and parameter inference performed here provide a unique resource for modeling the role market biosecurity can play, potentially bringing records of actual movements to bear on agricultural policy in ways not possible through existing [e.g. 53, 54] kernel-based models of disease spread [55]. Our study provides, for the first time, information on cattle movements that do not cross a state boundary. Although transportation of infectious cattle within a state would not immediately spark a regional epidemic, cattle movements at this scale could rapidly spread disease beyond the 10km control radius suggested in response to FMD detection within the US [1]. Our simple model of infection propagation stymied by market-based biosecurity measures highlights the tradeoffs involved in mitigating infection in the intensely-connected livestock industry of the US. In particular, we show that the efficiency of biosecurity measures at a market must be balanced against the ramifications of any delay in implementing movement bans.

## Acknowledgments

The authors want to acknowledge several undergraduate or graduate assistants who contributed to software development: Daniel Anderson, Adam Graves, Ching-Hao Hu, and Xinyang Jiang. We thank Nancy Robinson at the Livestock Marketing Association for sharing a member directory, and Centennial Livestock Auction of Fort Collins, CO for answering questions on cattle market practices. We also thank Jason E. Lombard (USDA-APHIS-VS) for several helpful conversations about livestock marketing practices. Funding for contributions by ITC and SB was provided through DHS Contract #HSHQDC-12-C-0014, with additional support from the RAPIDD Program of the Science & Technology Directorate, Department of Homeland Security, and the Fogarty International Center, National Institutes of Health.

## A Supplementary Data

Description of data release.

## B Supplementary Methods

Stan model code and elaboration of branching-process model.

## C Supplementary Figures

Individual market results as in Figure 2 and 4, and additional data on marketing arrangements.

## A Description of Data Release

**Note: The data release will coincide with publication of the article – the DOI given below will remain inactive until release.**

Data collected from livestock market websites, along with supporting data from sources attributed below, are available for download from the Bansal Lab Dataverse [1]. The data are separated into four tables that can be joined on one or more of the variables report, market and county. In addition, data on interstate flows [4] are available in electronic form at http://webarchives.cdlib.org/sw12j6951w/http://www.ers.usda.gov/Data/InterstateLivestockMovements/View.asp.

**receipts**

**report** (int, primary key) report identifier

**market** (int) foreign key linking to the distance table

**date** (char) date string as YYYY-MM-DD of the cattle auction

**receipts** (int) total number of cattle consigned

**mn** (char) USDA/AMS Market News (MN) report identifier

**mn** receipts (int) receipts from the corresponding MN report

The receipts for each cattle auction as reported by the auction market and as recorded in MN reports [2] for the same sale. Full MN reports are available online using the URL http://search.ams.usda.gov/mndms/{YYYY}/{MM}/{mn}.TXT with the substitutions indicated by curly braces.

**lots**

**lot** (int, primary key) lot identifier

**report** (int) foreign key linking to receipts

**head** (int) number of cattle

**county** (char) County code for the location of origin

The representative lots extracted from each market report with the consignor’s location geocoded to county, and the number of cattle consigned together (i.e. as a lot). The 

~~~
report
~~~

 identifier provides a link to additional information on each report in 

~~~
receipts
~~~

.

**distance**

**market** (int, primary key) market identifier

**county** (char, primary key) County codes used in this analysis

**distance** (decimal) miles along the great circle connecting centroids of this county and this market’s county

Distances are reproduced for convenience from [5]. The county containing a particular market is the one with zero distance.

**census**

**county** (char, primary key) County codes used in this analysis

**inv_num_cattle** (decimal) Inventory of cattle and calves

**inv_num_other** (decimal) Inventory of other cattle and claves

**inv_num_cows** (decimal) Inventory of cows and heifers that calved

**inv_num_beef_cows** (decimal) Inventory of beef cows

**inv_num_milk_cows** (decimal) Inventory of beef cows

**inv_num_on_feed** (decimal) Inventory of cattle on feed

**sale_num_cattle** (decimal) Sales by number of cattle

**sale_num_calves** (decimal) Sales by number of calves

**sale_num_500lbs** (decimal) Sales by number of cattle weighing over 500lbs

**sale_num_on_feed** (decimal) Sales by number of cattle on feed

These data are reproduced for convenience from [3] and are a selection of variables for the year 2012 census as seen in the county tables titled “Cattle and Calves – Inventory and Sales’ for each state (see http://www.agcensus.usda.gov/Publications/2012/ for PDF format and census methodology). The following relationships hold in principle,

~~~
inv_num_cattle = inv_num_cows + inv_num_other,
~~~

~~~
inv_num_cows = inv_num_beef_cows + inv_num_milk_cows, and sale_num_cattle = sale_num_calves + sale_num_500lbs.
~~~

Decimal values (rather than integer numbers of cattle) are possible due to the process of approximating missing data, which is described in the main text.

## B Supplementary Methods

### Branching Process Model

The random variable *Y* appearing in equation 4 represents the number of “children” for any “parent” in a branching process. Given a distribution on *Y*, it is possible to calculate the distribution on the size of the entire tree, although we only consider its first two moments. In the livestock disease scenario described in the main text, *Y* is the number of control areas which arise because a farm in a new area received cattle from an area that will also be designated a control area. Farms within a designated control area have all movements restricted, but there is a window between the first dangerous contact and restriction of all movements during which consignment of cattle at auction can induce further dangerous contacts.

We need several assumptions in determining a suitable distribution on *Y*, some to stand in for processes unrelated to any available data, such as the purchasing of auctioned cattle. We assume that the number of sales, *M*, at which operations from a given control area consign cattle during the time between the first dangerous contact by a farm within that area and the implementation of a movement ban has a Poisson distribution with mean *λ*. We denote by *N_i_* the number of cattle consigned at sale *i* of *M* by all the farms from an area soon to be controlled, and use market report data to estimate the distribution on *N_i_* as described in the main text. Finally, we assume that operations purchasing one or more of the *N_i_* cattle are distributed among *d* unique areas, so

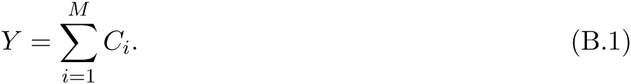

By independence among the *C_i_*, we have the following conditional expectation

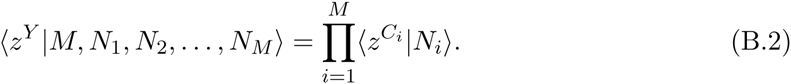

For the distribution on *C_i_*, we suppose a zero-inflation parameter *τ* gives the probability that the market hosting sale *i* quarantines all cattle from an area that will become controlled. Additionally, we suppose a splitting parameter *ν* that puts all buyers in the area where the cattle originate when *ν* = 0 and sends each animal to a unique area when *ν* = 1. In particular, we assume a binomial distribution on the range 0 to *N_i_* for the number of receiving areas when cattle are not quarantined, giving

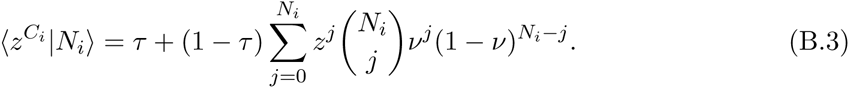

After averaging over the identically distributed *N_i_* to obtain 〈*z*^*C*^〉, and employing the generating function for the Poisson distribution,

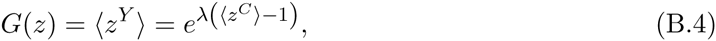

is the generating function for *Y*. From it, we obtain the first two factorial moments

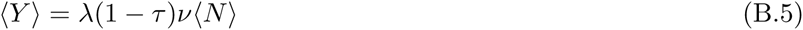

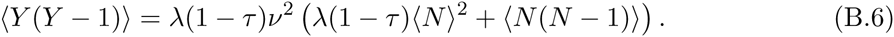

A generating function for the distribution on the total size of the branching process, or total number of control areas eventually established, is the obtained by application of the self-consistency relations described by Newman [1] for epidemics spreading on a random graph. We add the minor variation of keeping track of “generations”, so with random variable *X_n_* the size of a tree descending no more than *n* generations from a common ancestor, we impose the inter-generational consistency relation

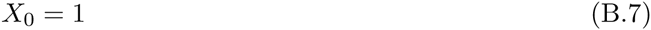

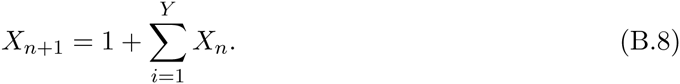

Because every *X_n_* in the sum is independent

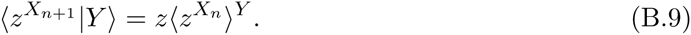

Denoting the generating function that has appeared on the right as *H_n_*(*z*), we obtain

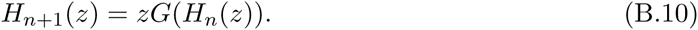

In theory, the number of control areas that must eventually be established depends on *H_n_*(*z)* as *n* → ∞, but the premise that *Y* does not tend to diminish over the course of an outbreak should not be taken too seriously. For an outbreak that is controlled within a small number of generations, however, the assumption of a stationary *Y* is a reasonable approximation.

### Stan Model Code

The following code in the Stan modelling language provides instructions for MCMC sampling of parameters affecting the likelihood on the number of cattle observed in representative lots consigned to a given market from different counties. Our likelihood function is the Pólya probability distribution parameterized by *α_j_* for *j* one of *c* counties (the subscript for market is dropped in this section because independent fits are performed for each market). The code implements a novel approach to calculating the likelihood for independent samples, i.e. the *n*_*j*,*i*_ head for each *i* of *s* reports from a focal market, drawn from a Pólya distribution, which would be naively calculated as:

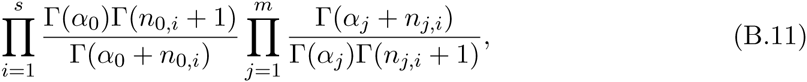

where 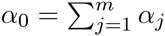 and 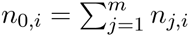 [2]. Calculating the log likelilood this way requires a routine for the log-gamma function, which must be called at least *m* + 1 times *for each sample*.

Calculating the log likelihood with an approach having constant complexity with respect to the number of samples is made possible by the identity

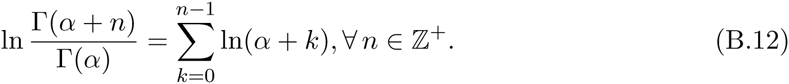

For the new approach, we first expand each log-gamma function according to this identity while using the Boolean function, *B*(·), to substitute for the summation limit:

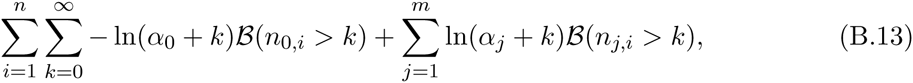

ignoring terms constant with respect to every *α_j_*. The sum across samples can now take place before parameter estimation begins, and the log likelihood is calculated as

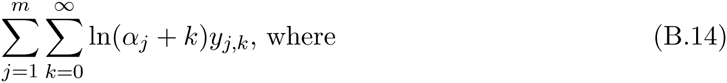

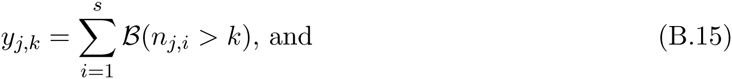

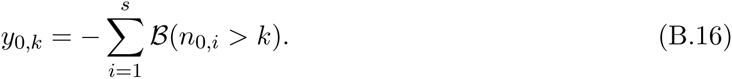

The double summation is implemented as a dot product in the code below, with sparse matrix *y*_·,·_. provided as a coordinate list (sparse COO storage) and with values of ln(*α_j_* + *k*) calculated as needed.

~~~
data {
~~~

~~~
/* Dependent variable */
~~~

~~~
int N; // number of non-zero values in sparse matrix
~~~

~~~
int M; // number of counties with observations
~~~

~~~
vector[N] y; // response value, in coordinate list (COO) sparse
~~~

~~~
vector[N] y_i; // matrix format, such that y[k] reports have more
~~~

~~~
int y_j[N]; // than y_i[k] animals from county y_j[k]
~~~

~~~
}
~~~

~~~
parameters {
~~~

~~~
vector<lower=0>[M] alpha;
~~~

~~~
}
~~~

~~~
model {
~~~

~~~
/* Prior */
~~~

~~~
alpha ~ cauchy(0, 5);
~~~

~~~
/* Likelihood */
~~~

~~~
{
~~~

~~~
vector[M + 1] x;
~~~

~~~
vector[N] z;
~~~

~~~
x[1:M] <- alpha;
~~~

~~~
x[M + 1] <- sum(alpha);
~~~

~~~
z <- log(x[y_j] + y_i);
~~~

~~~
increment_log_prob(dot_product(z, y));
~~~

~~~
}
~~~

~~~
}
~~~

The next code block specifies a multi-level, generalized linear model with log-link function and Pólya likelihood. The response variable is transformed as above into a sparse COO format, with a row for each livestock market and county pair. The hierarchical term adds a random intercept for each livestock market. The data under /⋆ Other ⋆/ provides indexing to correctly form and place 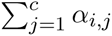 for each market *i* for use in the likelihood calculation.

~~~
data {
~~~

~~~
/* Dependent variable */
~~~

~~~
int N; // number of samples
~~~

~~~
int Q; // number of non-zero values in sparse sample matrix
~~~

~~~
vector[Q] y; // response value, in coordinate list (COO) sparse
~~~

~~~
int y_i[Q]; // matrix format, such that y[k] reports have more
~~~

~~~
vector[Q] y_j; // than y_j[k] animals from market:county y_i[k]
~~~

~~~
/* Categorical independent variables */
~~~

~~~
int K; // number of intercepts
~~~

~~~
int L; // number of variables
~~~

~~~
int C[L]; // position of last coefficient in each category
~~~

~~~
int M; // number of non-zero values in sparse design matrix for intercepts
~~~

~~~
vector[M] c_w; // design matrix in compressed row storage where c_w[i]
~~~

~~~
int c_v[M]; // is a value from column c_v[i], row j values begin
~~~

~~~
int c_u[N + 1]; // at position c_u[j] in c_w, and c_u[N + 1] = M.
~~~

~~~
/* Numerical independent variables */
~~~

~~~
int P; // number of slopes
~~~

~~~
matrix[N, P] x; // variables
~~~

~~~
/* Other */
~~~

~~~
int R_1;
~~~

~~~
int R_2;
~~~

~~~
int b_1[R_1];
~~~

~~~
int b_1_idx[R_1];
~~~

~~~
int b_2[R_2 + 1];
~~~

~~~
int b_2_idx[R_2];
~~~

~~~
}
~~~

~~~
parameters {
~~~

~~~
vector[C[L - 1]] alpha_1; // fixed intercepts
~~~

~~~
vector[C[L] - C[L - 1]] alpha_2_raw; // random intercepts
~~~

~~~
real<lower=0> sigma_2; // s.d. of random intercepts
~~~

~~~
vector[P] beta; // fixed slopes
~~~

~~~
}
~~~

~~~
model {
~~~

~~~
/* Prior */
~~~

~~~
append_row(alpha_1, beta) ~ normal(0, 100);
~~~

~~~
alpha_2_raw ~ normal(0, 1);
~~~

~~~
sigma_2 ~ cauchy(0, 5);
~~~

~~~
/* Likelihood */
~~~

~~~
{
~~~

~~~
vector[K] alpha;
~~~

~~~
vector[N] a;
~~~

~~~
vector[R_1 + R_2] b;
~~~

~~~
alpha <- append_row(alpha_1, alpha_2_raw * sigma_2);
~~~

~~~
a <- csr_matrix_times_vector(N, K, c_w, c_v, c_u, alpha) + x * beta;
~~~

~~~
a <- exp(a);
~~~

~~~
b[b_1_idx] <- a[b_1];
~~~

~~~
for (i in 1:R_2) {
~~~

~~~
b[b_2_idx[i]] <- sum(a[b_2[i]:(b_2[i+1] - 1)]);
~~~

~~~
}
~~~

~~~
increment_log_prob(dot_product(log(b[y_i] + y_j), y));
~~~

~~~
}
~~~

~~~
}
~~~

The last code block specifies a multi-level, generalized linear model with log-link function and negative binomial likelihood. The hierarchical term again provides a random intercept for each livestock market. Each record corresponds to one livestock auction with response variable equal to receipts minus the total number of cattle in representative lots.

~~~
data {
~~~

~~~
/* Dependent variable */
~~~

~~~
int N; // number of samples
~~~

~~~
int y[N]; // receipts - total of representative lots
~~~

~~~
/* Categorical independent variables */
~~~

~~~
int K; // number of intercepts
~~~

~~~
int L; // number of variables
~~~

~~~
int C[L]; // position of last coefficient per variable
~~~

~~~
int M; // number of non-zero values in design matrix
~~~

~~~
vector[M] d_w; // design matrix in compressed row storage where d_w[i]
~~~

~~~
int d_v[M]; // is a value from column d_v[i], row j values begin
~~~

~~~
int d_u[N + 1]; // at position d_u[j] in d_w, and d_u[N + 1] = M
~~~

~~~
/* Numerical independent variables */
~~~

~~~
int P; // number of slopes
~~~

~~~
matrix[N, P] x; // variables
~~~

~~~
}
~~~

~~~
parameters {
~~~

~~~
vector[C[L - 1]] alpha_1; // fixed intercepts
~~~

~~~
vector[C[L] - C[L - 1]] alpha_2_raw; // random intercepts
~~~

~~~
real<lower=0> sigma_2; // s.d. of random intercepts
~~~

~~~
vector[P] beta; // fixed slopes
~~~

~~~
real<lower=0> phi;
~~~

~~~
}
~~~

~~~
model {
~~~

~~~
/* Prior */
~~~

~~~
phi ~ cauchy(0, 5);
~~~

~~~
append_row(alpha_1, beta) ~ normal(0, 100);
~~~

~~~
alpha_2_raw ~ normal(0, 1);
~~~

~~~
sigma_2 ~ cauchy(0, 5);
~~~

~~~
/* Likelihood */
~~~

~~~
{
~~~

~~~
vector[N] log_mu;
~~~

~~~
vector[K] alpha;
~~~

~~~
alpha <- append_row(alpha_1, alpha_2_raw * sigma_2);
~~~

~~~
log_mu <- csr_matrix_times_vector(N, K, d_w, d_v, d_u, alpha) + x * beta;
~~~

~~~
y ~ neg_binomial_2_log(log_mu, phi);
~~~

~~~
}
~~~

~~~
}
~~~

## C Supplementary Figures

**Figure C.1:**
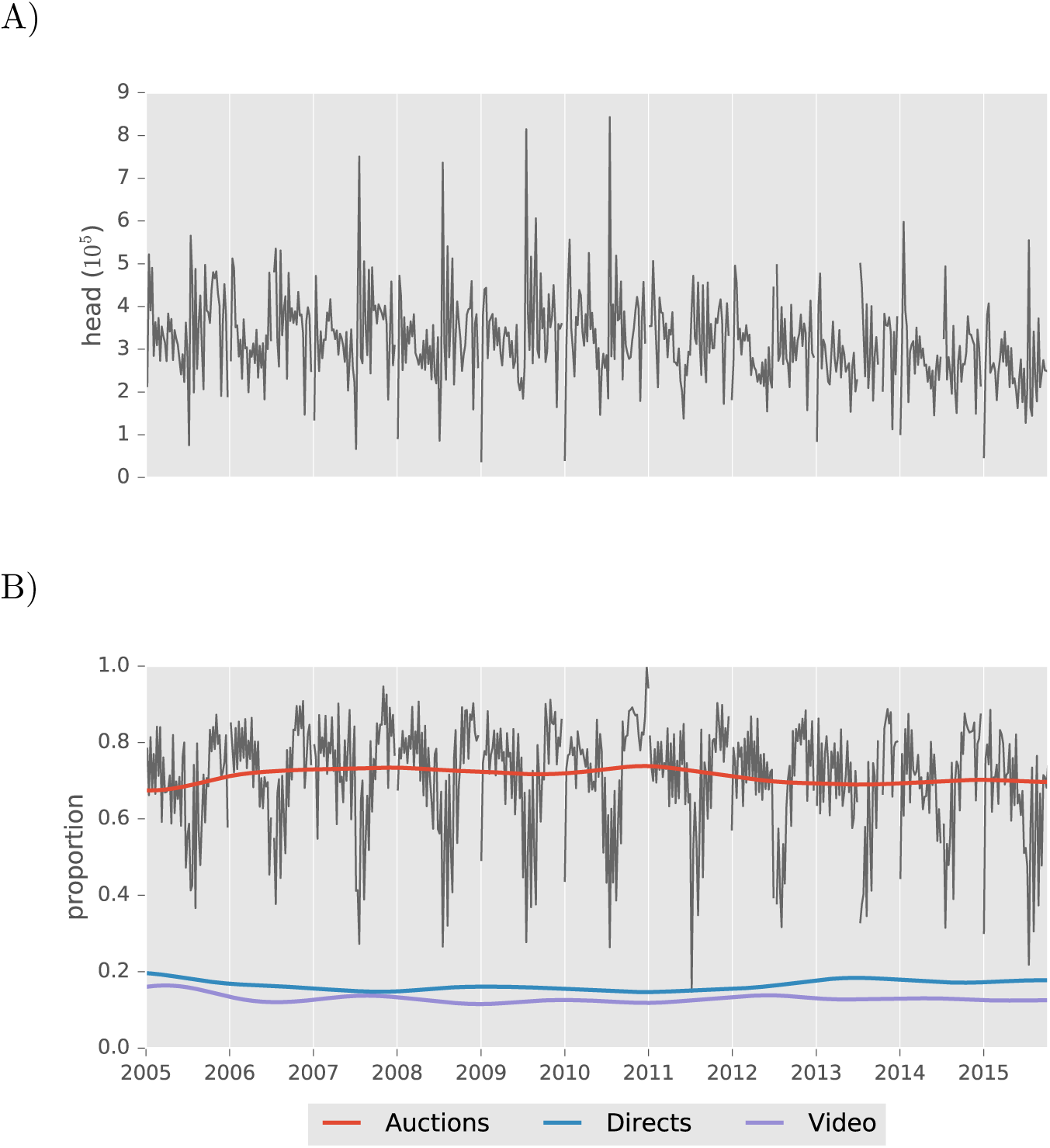
Feeder and stocker cattle, which are more epidemiologically relevant than cattle sold directly to meat packing operations, are predominantly sold through auction markets. A) Weekly volumes of feeder and stocker cattle sold. B) Proportion sold through auction markets (gray), and smoothed proportions for auction markets along with the next most common marketing arrangements. In contrast, the proportion of slaughter cattle sold through auction markets has declined in recent years [1]. National Feeder and Stocker Cattle Summary data were provided by USDA AMS.

**Figure C.2:**
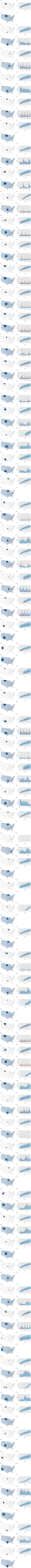
For each market, panels A, B and D correspond to analogous figures for a Montana market shown in the main text (Fig. 2A, Fig. 2B, and Fig. 4A, respectively). Panel C shows the posterior mean values of *e*^*x*_*i, j*_ *β_x_*^, binned according to the same boundaries separating quartiles in panel A. Even negligible values in Panel D fall into the first quartile, whereas estimates for *α*_*i*,*j*_ in panel A do not exist if county *j* was never observed to consign cattle at market *i*. Missing panels are due to receipts being unavailable for a market.

